# Antiviral Mx proteins have an ancient origin and widespread distribution among eukaryotes

**DOI:** 10.1101/2024.08.06.606855

**Authors:** Caroline A. Langley, Peter A. Dietzen, Michael Emerman, Jeannette L. Tenthorey, Harmit S. Malik

## Abstract

First identified in mammals, Mx proteins are potent antivirals against a broad swathe of viruses. Mx proteins arose within the Dynamin superfamily of proteins (DSP), mediating critical cellular processes, such as endocytosis and mitochondrial, plastid, and peroxisomal dynamics. And yet, the evolutionary origins of Mx proteins are poorly understood. Using a series of phylogenomic analyses with stepwise increments in taxonomic coverage, we show that Mx proteins predate the interferon signaling system in vertebrates. Our analyses find an ancient monophyletic DSP lineage in eukaryotes that groups vertebrate and invertebrate Mx proteins with previously undescribed fungal MxF proteins, the relatively uncharacterized plant and algal Dynamin 4A/4C proteins, and representatives from several early-branching eukaryotic lineages. Thus, Mx-like proteins date back close to the origin of Eukarya. Our phylogenetic analyses also reveal that host-encoded and NCLDV (nucleocytoplasmic large DNA viruses)-encoded DSPs are interspersed in four distinct DSP lineages, indicating recurrent viral theft of host DSPs. Our analyses thus reveal an ancient history of viral and antiviral functions encoded by the Dynamin superfamily in eukaryotes.

## Introduction

The vertebrate interferon (IFN) system acts as a first line of defense against viruses and other pathogens by inducing dozens of interferon-stimulated genes (ISGs) that create an antiviral environment. Functional IFN systems exist in early vertebrates, including bony fishes (1-3). Among the most rapidly and highly expressed ISGs upon interferon induction in human cells are Mx proteins, identified as antiviral proteins soon after the discovery of IFN (4-6). Although Mx proteins from different species have different antiviral specificities, they have an exceptionally broad range of activity. For example, the human MxA protein restricts diverse viruses, including Influenza A (7-9), Vesicular Stomatitis Virus (VSV) (8, 9), measles (10), and hepatitis B (11), whereas the human MxB paralog restricts retroviruses (12-14) and herpesviruses (15, 16). Most mammals have two *Mx* genes (17), although they were lost or pseudogenized in toothed whales (18). Birds have a single *Mx* gene, and fish encode up to seven *Mx* paralogs, which evolved by gene or genome duplication (1-3). Their well-documented presence in fishes and mammalian lineages suggested that the Mx proteins may have arisen coincident with the origin of the interferon system in the common ancestor of bony fishes and mammals. However, recent findings have revealed that ISGs such as STING and cGAS predate vertebrates (19-22). In addition, there have been two reports of Mx-like genes from invertebrate species (23, 24), suggesting an earlier origin.

Mx proteins are a member of the Dynamin superfamily of proteins (DSP), multi-domain GTPases that mediate many critical cellular processes within eukaryotic cells. Most DSPs localize to distinct cellular membranes, where they facilitate membrane remodeling. For example, Dynamin (or Dyn) proteins localize to the outer cellular membrane (25-33) and endosomes (34), whereas the Optic atrophy 1 (or Opa1) and Mitofusin (or Mfn) proteins act at mitochondrial membranes (35-41) alongside Dynamin-related proteins (or Drps) (42-46). In contrast to other studied DSPs, Mx proteins function independently of membranes (6, 47). Although Mx antiviral mechanisms are still poorly understood, one model proposes that they act by binding viral RNPs (ribonucleoproteins) and exerting a GTP hydrolysis-dependent power stroke to restrict virus replication (48), analogous to the power stroke exerted by Dynamin proteins on cellular membranes in the final step of endocytosis (49, 50).

Previous studies that deeply investigated the evolution and diversification of DSPs (51-53) included either very few or no Mx protein sequences in their analyses, leaving their evolutionary origins unclear. Conversely, studies on Mx evolution have focused exclusively on vertebrate or even mammalian Mx sequences (17, 54). Here, we analyzed the deep phylogenetic history of Mx in the context of DSPs using stepwise increments of eukaryotic phylogenetic coverage. We find unambiguous evidence that Mx-like proteins predate the birth of interferon in animals and are present within plants, fungi, and the majority of basally-branching eukaryotic lineages. Expanding our analyses to all eukaryotic DSPs, we reveal an ancient and ongoing history of lateral transfer between host genomes and nucleocytoplasmic large DNA viruses in four DSP lineages. Overall, our study reveals an ancient lineage of potentially antiviral Mx-like proteins in eukaryotes and an understudied potential arms race for dynamin-related functions between large double-stranded DNA viruses and their hosts.

## Results

### Mx predates the birth of interferon

To evaluate the evolutionary origins of Mx in the context of the broader Dynamin superfamily, we carried out BLAST searches on representative metazoan (animal) species with fully sequenced genomes using different human DSPs as queries.

Although many DSP genes undergo alternate splicing, we focused only on the longest isoform encoded by each DSP gene. We aligned all metazoan DSPs recovered with different query sequences using the MAFFT program (55). Because the GTPase domain is conserved across different DSPs, we manually extracted the GTPase domain from alignments of different DSPs and used these sequences to generate an all-metazoan DSP GTPase alignment, which was further trimmed manually. We subsequently used these sequence alignments to generate phylogenetic trees using FastTree (56, 57) (Figure 1A) or IQ-Tree (58, 59) (Supplementary Figure 1A).

**Figure 1.**
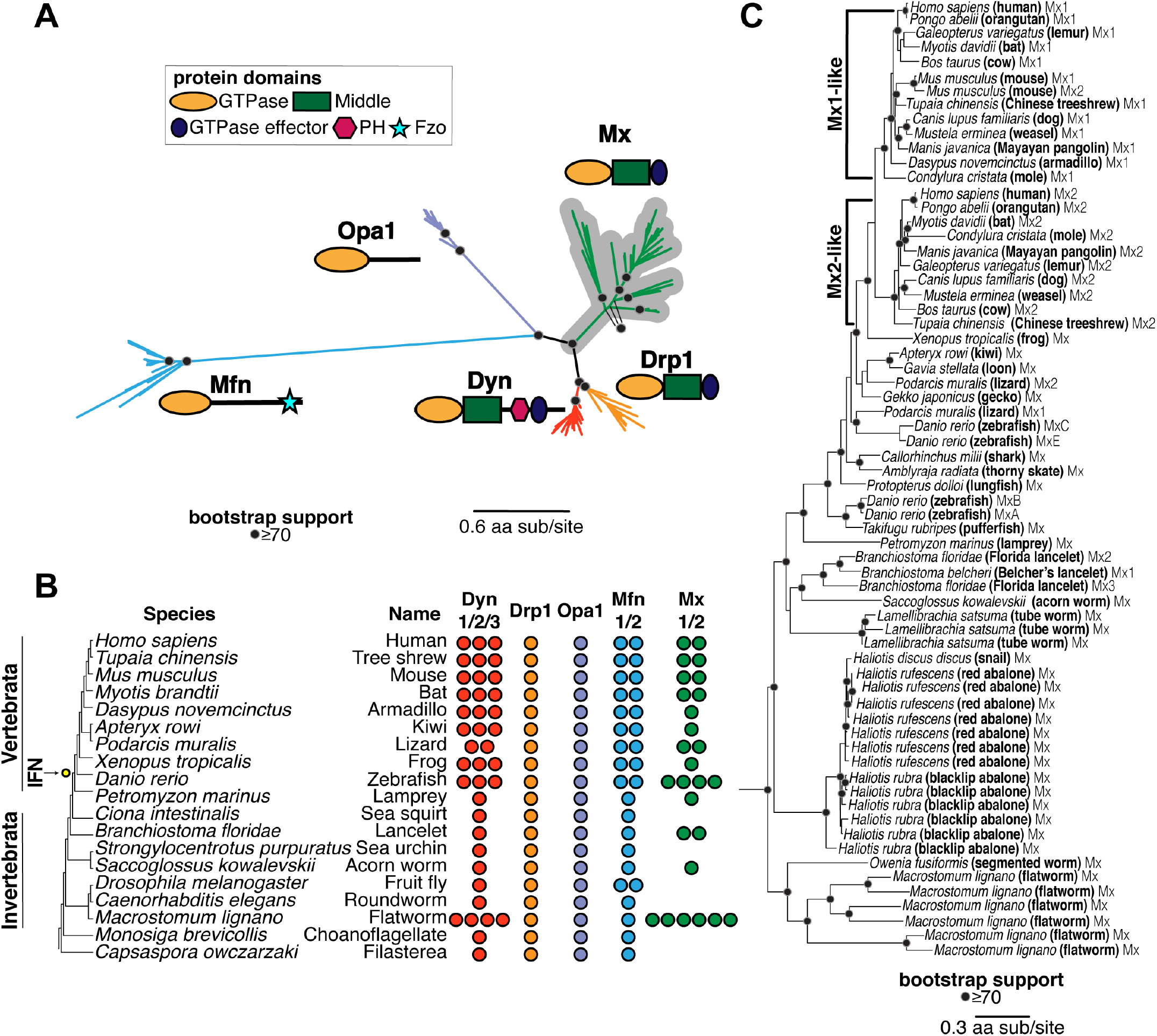
Evolutionary origin of Mx in animals predates the interferon signaling network. **(A)** Phylogenetic analyses of Dynamin-superfamily proteins (DSPs) based on their common GTPase domain in representative *Holozoa* (animals and their closest single-celled relatives) reveals five distinct DSP clades, consistent with previous analyses: Dyn, Drp1, Mfn, Opa1, and antiviral Mx (gray highlight). **(B)** Summary of localization of different DSPs in human cells. Dyn proteins localize to clathrin-coated pits on the plasma membrane, Drp proteins localize to mitochondria and peroxisomes, whereas Mfn and Opa1 proteins localize to mitochondrial membranes. In contrast, Mx proteins act independent of host membranes and localize to viral ribonucleoprotein (RNP) complexes in the cytoplasm (MxA) or proximal to the nuclear pore (MxB) or to the nucleoplasm (in other mammals, not shown). **(C)** Retention of different DSP clades in representative animals or their outgroup species (tree not drawn to scale). Drp1 and Opa1 are represented in a single copy in all animals and outgroup species, except for *S. arctica* (*Icthyosporea*), where we only recovered a partial Opa1 gene (indicated with a ‘?’). Mfn is also present in a single copy except in two cases. Mfn duplicated in bony vertebrates, giving rise to Mfn1 and Mfn2, coincident with the birth of the interferon system (IFN, yellow dot). Mfn independently duplicated in the lineage leading to *D. melanogaster*. Mx proteins are present in 1-4 copies in bony vertebrates and in 1-6 copies in some invertebrate species but have also been independently lost in several lineages. **(D)** Phylogenetic analysis of Mx-like proteins reveals a phylogenetic split between Mx1-like and Mx2-like genes in mammals and independent duplications in several fish lineages. We also find unambiguous evidence of basally-branching Mx-like genes in several invertebrate species, confirming that Mx genes have been vertically inherited in animals, followed by frequent subsequent loss and duplication events. (A), (D) Alignment generated by MAFFT and tree built using FastTree. Black dots indicate nodes with bootstrap support greater than 70% based on FastTree analyses (see methods); a scale bar indicates the level of amino acid divergence.

The resulting phylogeny (Figure 1A) reveals five distinct clades of DSPs with high bootstrap support, indicating high confidence in their phylogenetic relatedness. These five clades are consistent with the five DSP groups previously established in animal cells: Dyn, Drp1, Opa1, Mfn, and Mx proteins. Next, we analyzed the domain organization of the different homologs assigned to each of the five metazoan DSP clades using the NCBI conserved protein domain database (CDD) (60, 61). This analysis confirmed that all members of each DSP clade shared the characteristic domain differences previously used to distinguish DSPs (Figure 1A). For instance, a hallmark of canonical Dyn proteins is the Pleckstrin homology domain (PH) domain, which facilitates the recruitment of these proteins to the outer cellular membrane (62). Our analyses show that all metazoan sequences constituting the Dyn clade, and only sequences within this clade, encode a PH domain (Figure 1A). Similarly, all members of the Mfn clade encode the Fzo (fuzzy onion) domain, which appears to be restricted to this clade (Figure 1A). In contrast, the GTPase effector domain (GED) and Middle domains (that separate the GTPase from GED domains) are found in Dyn, Drp, and Mx proteins but not in Opa1 and Mfn proteins. Thus, the GTPase domain is the only universal domain common to all five DSP clades.

Proteins from these five DSP clades localize to distinct cellular compartments in human cells. Dyn proteins, which localize to the outer cellular membrane and other internal cellular membranes (25-34), phylogenetically group with Drp1 proteins, which localize to mitochondria and peroxisomes and are critical for organelle fission/fusion (42-46). Mx antiviral proteins, which have been shown to localize to the cytoplasm, the nucleoplasm, or the nuclear pore, form an outgroup lineage to the Dyn and Drp sister clades (6, 47). Opa1 proteins, which localize to the inner mitochondrial membrane (35-37), are an outgroup to the Mx, Drp, and Dyn clades. Much more basal branching is the Mfn clade, which encodes proteins that localize to the outer mitochondrial membrane (38-41).

Based on their phylogenetic groupings and protein-domain analysis (Figure 1A), we assigned all DSPs from representative metazoan species to each of the five distinct DSP clades (Figure 1C) to analyze instances of gene loss or duplication. We found that several DSP paralogs from the same clade are often present within the same species. For example, the human genome encodes three Dyn paralogs (Dyn1, Dyn2, Dyn3), two Mfn paralogs (Mfn1, Mfn2), and two Mx paralogs (MxA, MxB) but only one Drp1 and Opa1 (Figure 1C). A Mfn duplication in bony vertebrates gave rise to Mfn2, which modulates antiviral immunity (63, 64), whereas an independent Mfn duplication gave rise to the Marf and Fzo proteins in the *Drosophila* species. In addition to Dyn duplications in bony vertebrates, there are independent Dyn duplications in at least two invertebrate species: *Macrostonum lignano* (flatworm) and *Spaheoforma arctica* (Ic-thyosporea). In contrast, Drp1 and Opa1 appear to be encoded by single-copy genes in all metazoans, although we found only a partial Opa1 protein in *S. arctica* (indicated with a ‘?’ in Figure 1C).

These analyses also confirm the presence of Mx proteins in the vertebrate lineage and its absence in well-studied invertebrate models like *Caenorhabditis elegans* and *D. melanogaster*. However, we also find unambiguous evidence (based on bootstrap support and domain analysis above) of the presence of Mx orthologs in many invertebrate species, including *Branchistoma floridae* (lancelet), *Saccoglossus kowalevskii* (acorn worm), and *Macrostonmum lignano* (flatworm), which encodes at least six distinct Mx proteins. These findings suggest that Mx proteins arose in animals much earlier than the origin of the IFN gene network (in bony vertebrates). This ancient origin was followed by recurrent loss of Mx proteins from multiple invertebrate lineages and at least one lineage of mammals (18). This pattern of recurrent gene turnover (loss and duplication) is characteristic of host-virus evolutionary arms races between host and virus, as viral evolution renders some antiviral genes obsolete or imposes pressures on host genomes to expand antiviral functions via gene duplications (65).

To rule out the alternative possibility that the invertebrate Mx homologs might have resulted from horizontal gene transfers following their origins in vertebrates, we expanded our BLAST analyses to identify additional Mx proteins in animal genomes, using invertebrate Mx proteins as queries. We performed phylogenetic analyses using an alignment of the GTPase domain for all DSPs that unambiguously group within the Mx clade (Figure 1D). Our analyses recapitulate and extend findings from previous studies of Mx proteins in mammals (66, 67). We found two lineages of Mx proteins (Mx1 and Mx2) in mammals and Mx representatives from bird, amphibian, shark, and fish lineages. In invertebrates, we identified Mx homologs in *Lamellibrachia satsuma* (tube worm), multiple *Haliotis* species (snail, abalone), and *Owenia fusiformis* (segmented worm) (Figure 1D, Supplementary Figure 1B) in addition to the previously identified lancelet, acorn worm, and flatworm lineages (Figure 1B, Figure 1C). Most importantly, the topology of the Mx tree largely mirrors the species tree, consistent with the early origin of Mx proteins in the animal phylogeny. Together, these analyses show that animal Mx proteins are more ancient than the interferon system in vertebrates, have largely been subject to vertical inheritance, and have undergone several lineage-specific gene duplications and losses.

### Phylogeny of animal, fungal, and plant DSPs reveals ancient Mx orthologs

Based on our finding that the Mx clade arose early in the origins of animals, we wanted to extend our analyses of potential Mx origins to two additional lineages – fungi (a sister lineage to animals) and plants – in which DSPs have also been well-studied. Using the same approach of iterative BLAST searches of representative animal, fungal, and plant genomes using different DSP queries, we carried out phylogenetic analyses of all DSPs recovered from these genomes based on their common GTPase domain with FastTree (Figure 2A) and IQ-Tree (Supplementary Figure 2A). We also analyzed their domain architecture using the CDD (Figure 2A). These analyses revealed fungal and plant orthologs of animal Mx proteins. For example, animal Mx proteins unambiguously group with uncharacterized DSPs in some fungi, which we rename MxF (for Mx-like proteins from Fungi). For example, *Aspergillus fumigatus* encodes five MxF proteins, *Agaricus bisporus* encodes three, and *Batrachochytrium dendrobatidis* encodes one (Figure 2B). In contrast, many different fungal lineages encode no MxF proteins at all *(e*.*g*., *Saccharomyces cerevisiae, Ustilago maydis, Piromyces sp. E2, Conidiobolus coronatus, and Allomyces macrogynus*) (Figure 2B). This extreme dynamism in copy number is highly reminiscent of the gene loss/ expansion seen in Mx genes in animal genomes but also explains why previous studies failed to identify MxF genes in fungi or misclassified them as Dynamin proteins. Despite their heterogeneous presence, MxF proteins are found in most major clades of fungi (Figure 2C), including *Ascomycota, Basidiomycota, Chytrid*, and *Mucoromycota* (indicated with ‘A’, ‘B’, ‘C’, and ‘M’ in Figure 2C), and, as well as *Aphelida*, which are believed to the sister lineage to true fungi (68, 69).

**Figure 2.**
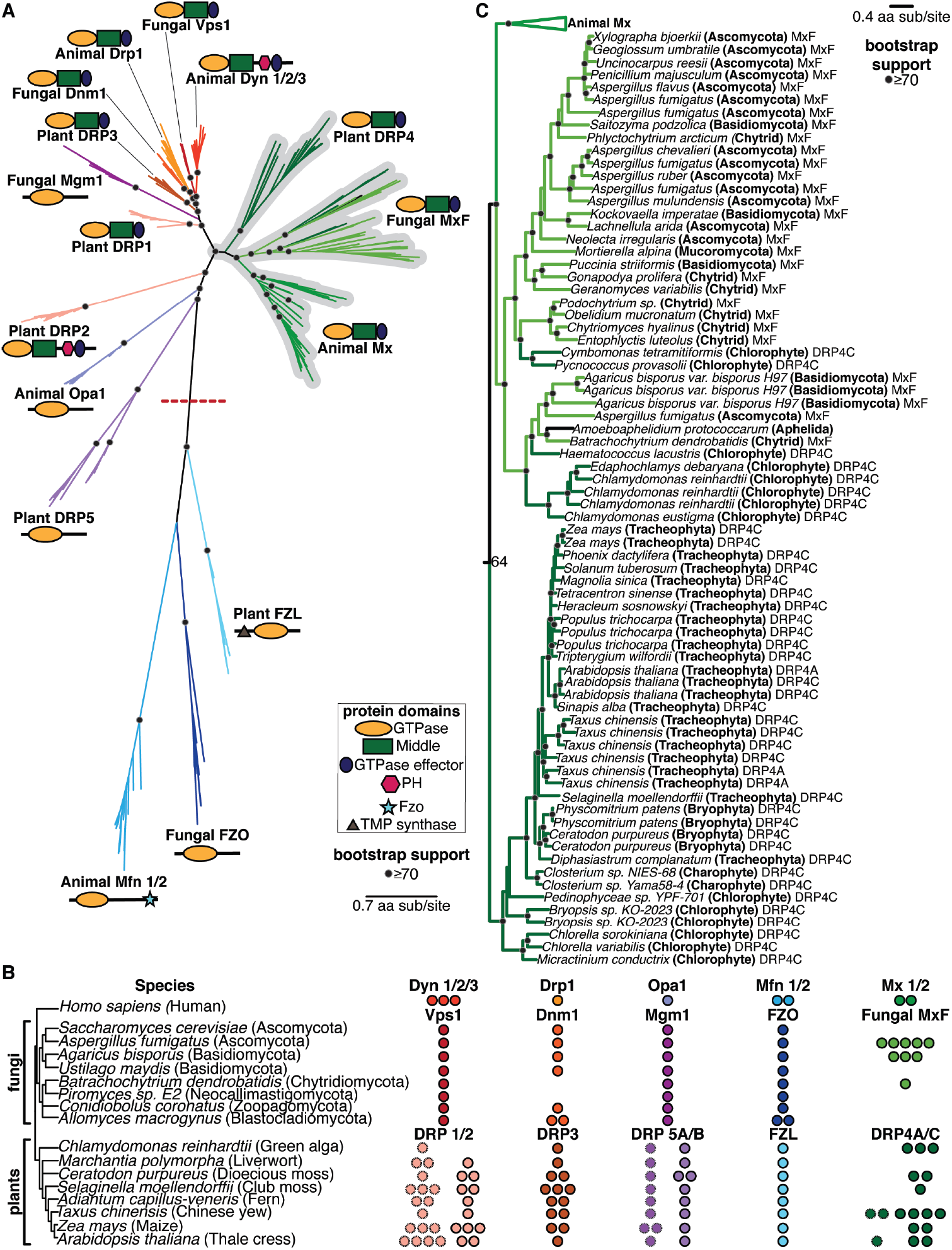
Animal DSP clades, including Mx proteins, in fungal and plant genomes. **(A)** Phylogenetic analysis of animal, fungal, and plant DSPs based on their common GTPase domain reveals broad groupings. Animal Dyn proteins group with fungal Vps1 proteins but not with plant DRP1 or DRP2 proteins, even though only Animal Dyn and plant DRP2 proteins share a C-terminal PH domain. Animal Drp proteins group functionally with fungal Dnm1 and plant DRP3 proteins. Although Fungal Mgm1 is considered the functional equivalent of animal Opa1 and plant DRP5A/5B proteins, our phylogenetic analyses suggest that they are not true orthologs. In contrast, animal Mx proteins appear unambiguously orthologous to uncharacterized fungal MxF proteins and plant DRP4 proteins. Finally, the divergent animal Mfn lineage groups with fungal FZO and plant FZL proteins, which are very divergent from the rest of the DSPs. **(B)** Representation of various DSP classes in representative fungal and plant (and green algae) genomes. Among fungi, Vps1, Mgm1, and FZO are encoded by single-copy genes (except for an *FZO* duplication in *A. macrogynus*). *Mx*-like genes vary from zero to six copies (in *A. fumigatus*). Among plants, FZL is encoded by a single copy gene in all representative algae and plants. DRP5A and DRP5B are mostly also encoded by single-copy genes in plants (except for a DRP5A duplication in maize and a DRP5B duplication in a moss species). DRP1 (hatched outline) varies from one to four copies in plants, whereas DRP2 (solid outline) is present in one to three copies (and absent in green algae). DRP3 varies from one to three copies in all algae and plants. Finally, DRP4C is present from zero to three copies. In addition to DRP4C (solid outline), many plants also encode shorter DRP4A proteins (hatched outline). **(C)** Phylogenetic analysis reveals a monophyletic lineage consisting of animal Mx, fungal MxF (Mx-like proteins from Fungi), and plant DRP4 proteins. MxF proteins are represented in a single lineage found in a variety of fungal lineages, including Ascomycota (A), Basidiomycota (B), Chytrids (C), and Mucoromycota (M), as well as Aphelia, which are a pseudo-fungi-like sister lineage to Fungi. Most Plant DRP4C proteins from various lineages of plants – *Chlorophytes, Charophytes, Bryophytes, and Tracheophytes* are also found in a single lineage, with full-length DRP4C proteins occasionally interspersed with shorter DRP4A proteins. However, some *Chlorophyte* DRP4C proteins group with fungal MxF proteins rather than other DRP4C, which may indicate at least two MxF fungal-to-algal horizontal transfer events. (A), (C) Alignment generated by MAFFT and tree built using FastTree. Black dots indicate nodes with bootstrap support greater than 70% based on FastTree analyses (see methods); a scale bar indicates the level of amino acid divergence.

Consistent with a previous proposal (70), we find that plant DRP4 proteins are orthologous to animal Mx and fungal MxF proteins (Figure 2A, Figure 2C). Plants encode full-length DRP4C proteins, which resemble animal Mx proteins in length and domain architecture, and much shorter DRP4A proteins, which often comprise only a GTPase domain with a truncated stalk domain. We find that DRP4A and DRP4C genes from the same species are often more closely related to each other than to orthologs in different plant species, suggesting that DRP4A genes might have arisen independently multiple times in plant evolution from full-length DRP4C proteins. Since we never find plant genomes that only encode the DRP4A proteins, we speculate that shorter DRP4A genes might have recurrently arisen to regulate the activity of full-length DRP4C proteins. Detailed phylogenetic analyses (Figure 2C, Supplementary Figure 2B) reveal that DRP4 proteins are widespread in lineages of green algae and plants and are present in many lineages, including *Chlorophytes, Charophytes* (also a green algal lineage), *Bryophytes* (non-vascular plants), *and Tracheophytes* (vascular plants, which include ferns, gymnosperms, and angiosperms). Our analyses also revealed two obvious instances of potential horizontal gene transfer (HGT) from fungi to chlorophytes (Figure 2C, Supplementary Figure 2B). The first instance occurred (from a chytrid MxF) into the ancestor of two Chlorophytes – *Pycnococcus provasolii* and *Cymbomonas tetramitiformis* – while the second event occurred into *Haematococcus lacustris, Edaphochlamys debaryana*, and *Chlamydomonas reinhardtii*. These are indicated as DRP4C proteins in our phylogeny (Figure 2C, Supplementary Figure 2B) but are more likely MxF proteins. Like animal Mx and fungal MxF proteins, we identify extremely dynamic gene turnover within plant DRP4 proteins, consistent with their engagement in evolutionary arms races with viruses as *bona fide* antiviral proteins.

We conclude that the Mx lineage is much more ancient than previously believed and includes representatives of animals, fungi, and plants.

### Phylogenetic relationships between other animal, fungal, and plant DSPs

Our analyses also reveal insights into and clarify the phylogenetic relationships between the other DSPs. For example, the ‘Drp’ grouping of animal Drp proteins, fungal Dnm1 (dynamin-related GTPase), and plant DRP3 (Figure 2A), is consistent with their localization and function in mitochondria and peroxisomes (70-75). Dnm1 is encoded by a single copy gene in representative fungi, except for a duplication in *Allomyces macrogynus* and a loss in both *Batrachochytrium dendrobatidis* and *Piromyces sp. E2*, whereas *DRP3* genes are present in 1-3 copies in all representative plant species (Figure 2B). The apparent loss of Dnm1 in some fungal species is unexpected, given their essential roles in many organisms; this might suggest functional redundancy between different DSP clades.

In contrast, the ‘Dyn’ grouping is more puzzling at first glance. Fungal Vps1 (vacuolar protein sorting) proteins, present in a single copy in most fungi, are closest in sequence and considered the functional equivalent of animal Dyns (76, 77) (Figure 2A). Vps1 proteins are implicated in vacuolar fusion (78-80), membrane scission (79-82), and peroxisomal partitioning (73, 83-85). Most plants encode 1-3 copies of two Dyn-like proteins, DRP1 and DRP2, which play a role in clathrin-mediated endocytosis (70, 86-91) and at the cell plate during cytokinesis (92, 93). And yet, neither fungal Vps1 nor plant DRP1 proteins encode a pleckstrin homology (PH) domain, a defining characteristic of animal Dyn proteins. In contrast, despite being highly divergent from animal Dyn proteins, plant DRP2 proteins (Figure 2A) encode a PH domain. We addressed this apparent contradiction by making separate phylogenies of all Dyn, Drp, and Mx proteins based either on their shared GTPase domains (as before, Figure 2A) or their shared Middle and GED domains (Supplementary Figure 3). Based on the Middle-GED domain phylogeny, we find that plant DRP1 and DRP2 proteins are sister lineages (Supplementary Figure 3), even though the DRP2 appears to be much more divergent than DRP1 in the GTPase phylogeny (Figure 2A). We posit that an ancestral plant DRP1/2 protein, encoding a PH domain, duplicated to give rise to DRP1, which lost the PH domain, and DRP2, which likely acquired a divergent GTPase domain via recombination (Supplementary Figure 3). An alternative possibility is that plant DRP1 GTPase domains evolved more rapidly than other DSPs, leading to their divergent placement in the GTPase phylogeny. Although green algae *Chlamydomonas reinhardtii* only encodes DRP1, most other plants encode 1-3 copies of DRP1 and DRP2 (Figure 2B, Supplementary Figure 3).

Fungal Mgm1, which shares the overall domain architecture as animal Opa1, maintains mitochondrial ultrastructure and morphology and regulates mitochondrial fusion like Opa1 (94-96). And yet, the phylogenetic grouping of fungal Mgm1 and animal Opa1 proteins is not very strong (Figure 2A). Like Opa1 in animals, Mgm1 is encoded in a single copy in most fungi, while most plant genomes encode 1-2 copies of each of the DRP5A and DRP5B paralogs (Figure 2B). Based on their similar structure, plant DRP5 proteins should be excellent candidates for functional equivalents of animal Opa1. However, unlike animal Opa1 proteins, which exclusively function in mitochondria, plant DRP5 proteins function in cytokinesis (75), chloroplast and peroxisome division (97, 98), and mitochondrial morphogenesis/division (99). Thus, the ‘Opa1’ grouping, consisting of animal Opa1, fungal Mgm1, and plant DRP5 proteins, does not show strong evidence of monophyly (Figure 2A) or functional similarity.

The animal Mfn, fungal FZO, and plant FZL proteins, which are mostly encoded by single-copy genes (Figure 2B), group together to the exclusion of the rest of the DSPs (Figure 2A). Animal Mfn and fungal Fzo proteins mediate the interaction between mitochondrial outer membranes to drive mitochondrial fusion (100, 101), whereas plant FZL proteins localize to chloroplasts and function in thylakoid organization (102). Thus, it is unclear whether this grouping reflects true orthology or is simply a result of their high divergence from the rest of the DSPs.

### Deep evolutionary origins of the Mx-like DSPs in eukaryotes

We expanded our survey of Mx-like and other DSP proteins beyond animals, fungi, and plants to diverse basally-branching eukaryotes and eukaryotic viruses (Figure 3A, Supplementary Figure 4A). We did not include Mfn, FZO, and FZL proteins in this analysis since they are quite divergent from the remainder of the eukaryotic DSPs. Our survey identified basally branching eukaryotic representatives in clades that were already well-established (Figure 3A). For example, we found several basal branching eukaryotes, including Stramenopiles, Alveolates, Rhizaria, Amoebazoa, Discoba, and Haptopytes, in the Dyn and Drp clades, suggesting that the Dyn/ Drp clade was already present as a fully specialized, distinct DSP clade in the last eukaryotic common ancestor (LECA).

**Figure 3.**
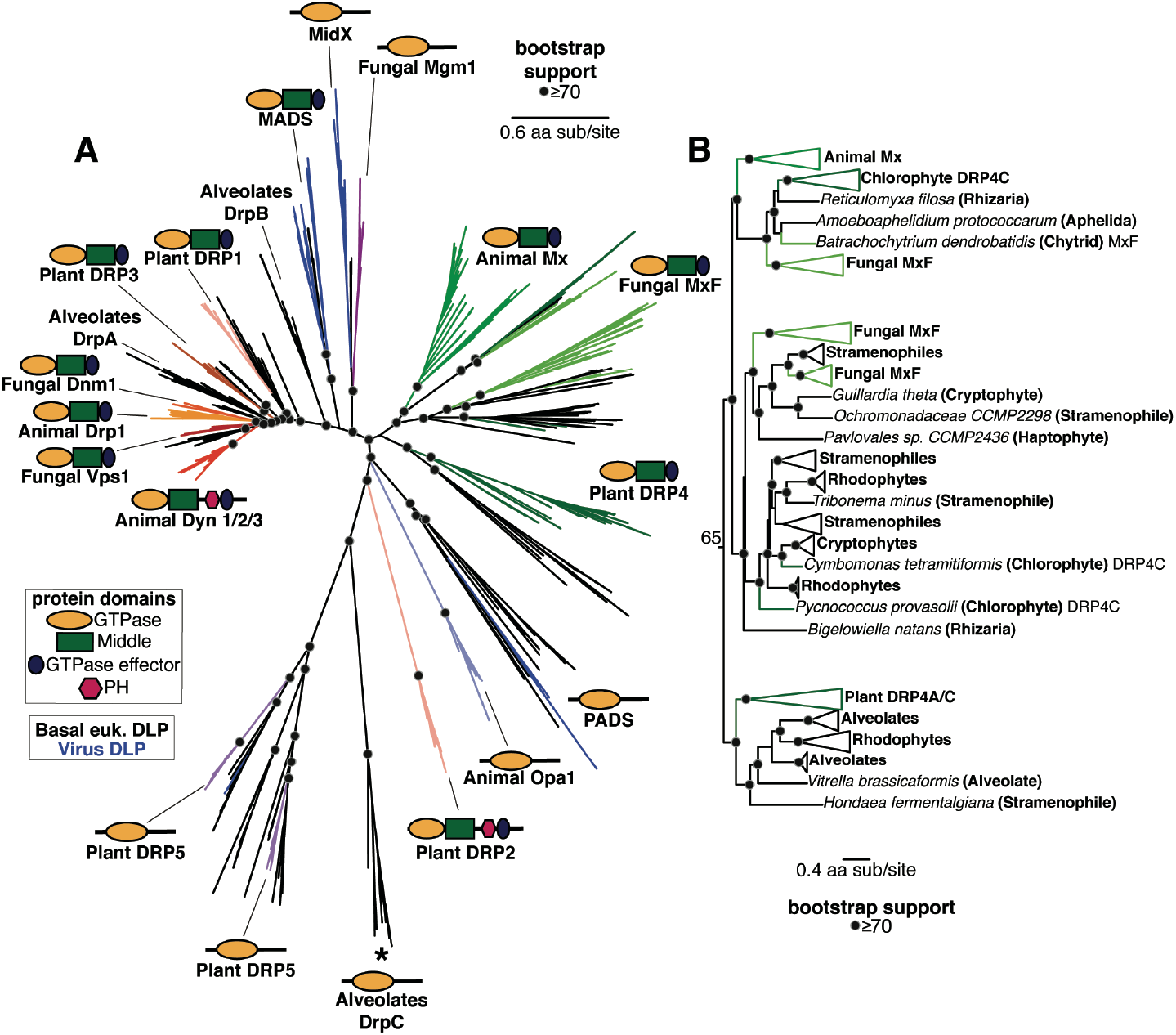
Phylogenetic analysis of DSPs in eukaryotes reveals an ancient Mx lineage. **(A) Phylogenetic analysis of the GTPase domain from all eukaryotic DSPs (except for the Mfn/FZO, FZB, and FZL clades) reveals an ancient Dyn/Drp clade that includes** Animal Drp, Plant DRP3, and Fungal Vsp1 along with fungal Dnm1, alveolate DrpA and DrpB, Plant DRP1, and DSPs from several basal branching eukaryotes (shown with black branches). Branching outside the Dyn/Drp cades are the MADS and MidX/ Fungal Mgm1 clades; blue branches indicate DSPs found encoded in viral genomes. Branching next is a single monophyletic clade of Mx-like DSPs (described in more detail in (B)) and the newly discovered PADS lineage. Finally, we find a grouping of animal OPA1, Plant DRP5 (which also contains Stramenopile, Amoebazoa, and a single virus DSP), and Alveolate DrpC. Although Plant DRP2 also groups with this final grouping, we believe its correct phylogenetic position (based on the Middle and GED domains) is as a sister to Plant DRP1 (Suppl Figure 1). **(B)** Our phylogenetic analysis delineates three deeply branching lineages of Mx-like proteins in eukaryotes. The first of these consists of representatives from animals, fungi, algae, and Rhizaria. The second lineage consists of representatives from fungi, Stramenopiles, Haptophytes, Cryptophytes, Chlorophytes, and Rhizaria. Finally, the third deep lineage consists of Mx-like sequences from plants, alveolates, rhodophytes, and Stramenopiles. (A), (B) Alignment generated by MAFFT and tree built using FastTree. Black dots indicate nodes with bootstrap support greater than 70% based on FastTree analyses (see methods); a scale bar indicates the level of amino acid divergence.

Our analyses also revealed Mx-like proteins in basally-branching eukaryotes (Figure 3A, Figure 3B). The Mx-like proteins we have identified branch in three deep lineages (Figure 3B, Supplementary Figure 4B). The first lineage comprises animal Mx proteins, fungal MxF proteins (including *Chytrids* and *Aphelida*), chlorophyte (green algae) DRP4C, and Mx-like representatives from Rhizaria, unicellular eukaryotes that are part of the TSAR (Telonemia-Stramenopiles-Alveolates-Rhizaria) supergroup of eukaryotes (103, 104). The second lineage consists of Mx-like proteins from Stramenophiles (also referred to as Heterokonts), Cryptophytes (a group of divergent plastid-bearing algae, also referred to as Cryptomonads), Haptophytes (a distinct divergent group of algae), Rhodophytes (red algae), and Rhizaria. The third deep lineage of Mx-like proteins consists of Plant DRP4C, and Mx-like representatives from Alveolates, Rhodophytes, and Stramenopiles. Thus, Mx-like proteins are found in representatives of the majority of the early-branching supergroups of extant eukaryotes, including TSAR (e.g., Stramenopiles, Alveolates), Haptists (*e*.*g*., Haptophytes), Archaeplastida (*e*.*g*., red algae, green algae, plants), Cryptista (*e*.*g*., Cryptophytes), and Amorphea (*e*.*g*., animals, fungi). Based on this representation in early branching eukaryotic supergroups, we infer that Mx proteins arose close to or shortly following LECA, much earlier than previously suspected.

Our analyses also identify eukaryotic DSP lineages that were either previously unidentified (PADS) or only recently identified (MADS, MidX) (53), which provide additional clarity about the phylogenetic relationships between different DSPs. For example, consistent with findings from a recent study (53), we also find the fungal Mgm1 clade to be more closely related to the newly identified MidX clade with high bootstrap support rather than the rest of the Dyn/Drp homologs. Thus, fungal Mgm1 proteins are phylogenetically not as closely related to the lineage that includes animal OPA1, alveolate DrpC, and plant DRP5 proteins. Similarly, the plant DRP2 GTPase domain appears to be more closely related to the MADS lineage of DSPs, consistent with our previous hypothesis that the DRP2 lineage may have swapped its original DRP1-like GTPase domain with a more divergent GTPase (Supplementary Figure 3). Finally, we uncovered four distinct DSP clades that contain genes from both eukaryotes and viruses, strongly indicative of horizontal gene transfer (Figure 4). For example, the MADS clade (previously described as “Clade D” (53)) is found in haptophytes (*Emiliana huxleyi, Chrysochromulina tobinii*, and *Dia-cronema lutheri*) as well as interspersed lineages of *Nucleo-cytoviricota* (nucleocytoplasmic large DNA viruses, or NCLDV) (blue lineages, Figure 4). Similarly, the PADS lineage consists of DSPs from haptophytes, chlorophytes (green algae), oomy-cetes (which are part of the Stramenopiles), and NCLDVs (Figure 4). Previous studies uncovered a large insertion from an ancient giant virus in oomycete genomes (105), but we found no evidence that the oomycete PADS sequence originates from this insertion. Our analyses also uncovered MidX sequences from NCLDV DSPs (Figure 4). Although we did not recover any MidX sequences from eukaryotic host genomes based on our analyses of the well-curated non-redundant database in NCBI, metagenomic data has revealed additional host MidX sequences in a recent study (53). Finally, we also recovered a DRP5-like DSP sequence from a *Clandestinovirus* NCLDV nestled within the DRP5 phylogeny from *Amoebozoa* (106). In nearly all cases, the viral DSPs have perfectly preserved all the catalytic residues that would indicate retention of GTPase activity (Figure 4B) (107, 108). Thus, PADS, MADS, MidX, and DRP5 DSPs represent lineages that have undergone HGT between eukaryotic host cells and their resident large viruses (Figure 4A).

**Figure 4.**
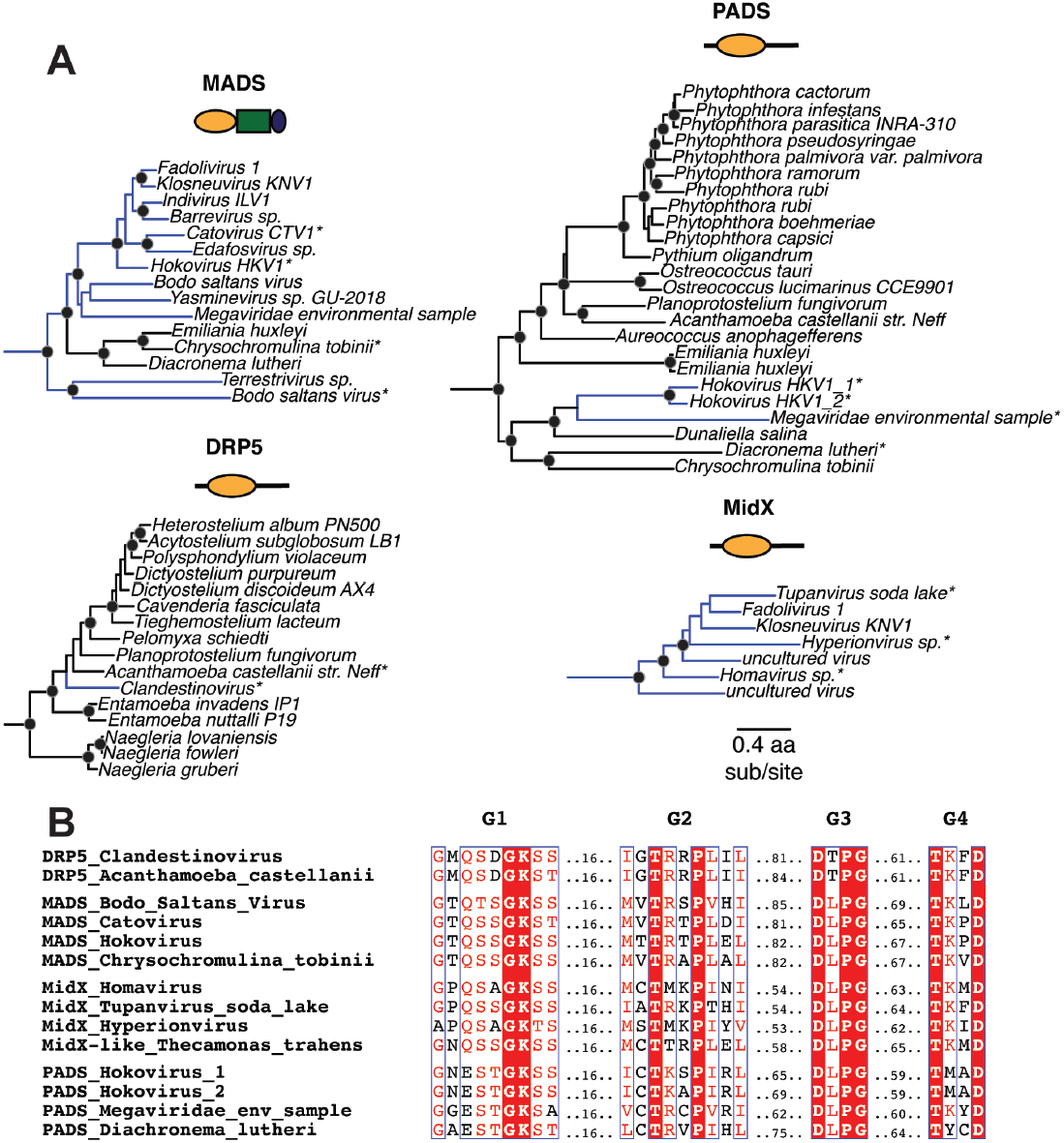
Viruses acquired host DSPs in at least four lineages. **(A)** We find that at least 4 major DSP clades contain viral DSP sequences, suggesting an ancient history of HGT between host and virus. MADS mostly contains NCLDV sequences, with three basal branching eukaryotic DSPs as outgroups. PADS consists of a few NCLDV sequences that serve as an outgroup to a diverse clade of basal branching eukaryotes. Our analysis only uncovered viral MidX DSPs. However, previous studies also un-covered eukaryotic representatives from metagenomic data (52). Finally, nestled among DRP5 sequences from *Ameoebozoa* is a single NCLDV DSP. Alignment generated by MAFFT and tree built using FastTree. Black dots indicate nodes with bootstrap support greater than 70% based on FastTree analyses (see methods); the scale bar indicates the level of amino acid divergence for all four clades shown here; asterisks indicate samples used in B. **(B)** Alignment of G Box motifs in viral DSPs indicate preservation of catalytic motifs associated with GTP hydrolysis in viral DSP sequences.

Thus, in addition to the deep evolutionary origin of Mx proteins in eukaryotes, with recurrent gene turnover — a hallmark of antiviral function — we find that large DNA viruses have also recurrently coopted DSP proteins. Our phylogenetic results suggest that the critical and yet adaptable functionality of DSP proteins has placed them center-stage in the evolutionary battles between hosts and viruses.

## Discussion

Given their essential roles in cellular remodeling, there has been considerable interest in the evolutionary and functional diversification of Dynamin-like proteins in eukaryotes. Previous analyses have taken advantage of increased information about cellular localization, structural information, and burgeoning sequencing databases to propose schemes for how this critical gene family arose and diversified in eukaryotes (17, 29, 53, 62). We build upon these earlier studies to deeply investigate the origins of the unusual Mx antiviral DSPs in eukaryotes, which were poorly understood. We find that the Mx lineage of DSPs is much more ancient than initially believed. Some earlier reports had suggested that its evolutionary origins coincided with the birth of the interferon system in bony vertebrates (17, 54). Instead, our analyses reveal distinct Mx-like lineages in many animals (including invertebrates), fungi, plants, and basal-branching eukaryotic supergroups.

It is unclear whether all of these Mx-like lineages also perform antiviral roles, just like they do in vertebrates, although there are some suggestions this might be the case. For example, some plant DRP4C proteins have been suggested to participate in antiviral function (62, 70, 75). Moreover, if Mx-related proteins were instead performing some necessary cellular membrane remodeling function in non-animal species, we would expect much less genetic turnover; instead, we find both rampant gene amplification (in animals, plants, and fungi) as well as several instances of complete gene loss (in animals and fungi). This gene turnover is more consistent with antiviral rather than essential cellular housekeeping functions. Our findings suggest that Dynamin specialization for antiviral function occurred early in eukaryotic evolution. The origins of other interferon-induced antiviral genes, such as cGAS, STING, and Viperin, lie in bacteria (20, 21, 109). However, our analyses could not find a bacterial or archaeal ortholog to eukaryotic Mx-like genes, even though at least one bacterial Dynamin-like protein participates in antiviral defense (110).

Our analyses reveal more details about early events of Dynamin specialization, including novel DSP lineages whose cellular functions are still unknown. For example, a recent study described and characterized the MidX lineage, which can remodel mitochondrial membrane topology from within the matrix (53), unlike Opa1 and Mfn, which remodel mitochondrial membranes from the outer membrane and intermembrane space. In addition to MidX and MADS, we identify a novel PADS lineage. Our findings indicate that much DSP diversification in extant eukaryotes must have already occurred in early eukaryotic evolution. These findings challenge previous classifications of ‘modern’ versus ‘ancient’ dynamins; these earlier classifications might have been influenced by the heavy over-representation of well-studied model systems, ***i*.*e***., animals, fungi, and plants, at that time.

Another common theme across the DSPs is the recurrence of horizontal gene transfers (HGTs), including the fungal-toalgal transfer of Mx-like genes (Figure 2C), which implies an exchange of antiviral function in some cases. Understanding the functional consequences of these inferred HGTs might provide further insight into the function of the non-animal Mx proteins. Most intriguing among these HGT events is the interspersing of eukaryotic and viral DSPs in at least four distinct DSP lineages – MidX, MADS, PADS, and DRP5 – which suggests an ancient, ongoing history of HGT of DSPs between multiple eukaryotic host lineages and *Nucleocytoviricota* (NCLDV). NCLDV are a family of double-stranded DNA viruses, including *Mimiviridae, Poxviridae, Asfarviridae, Iridoviridae*, and *Phycodnaviridae*, typified by large genomes and viral particle sizes (111). These viruses infect many eukaryotes, including algae, amoeba, and animals (112-114). Among viruses, only NCLDVs encode Dynamin-like proteins, suggesting some unique aspect of their cell biology may require Dynamin-like function. Unlike other lineages of double-stranded DNA viruses, NCLDV replication and assembly take place almost entirely within the host cytoplasm, with no discernible steps taking place in the host nucleus. The discovery of novel and complex membrane remodeling in *Molliviridae, Mimiviridae*, and *Poxviridae* highlights specific cell biological requirements for assembling large viral particles of NCLDVs in the host cytoplasm (115-119). Thus far, host Dynamin proteins have not been directly implicated in these processes. However, the study of cell biology of most NCLDVs is still in its infancy. Our work and others (53) suggest that NCLDV assembly of new envelope-bound virions may require or be enhanced by Dynamin-like activity, which may have spurred at least some NCLDVs to acquire and repurpose host DSPs for their function. An alternative model is that NCLDVs acquire host DSPs specifically to interfere with host DSP-mediated processes by a ‘dominant-negative’ model. However, if this were the case, we would not expect universal retention of all features required for GTPase catalytic activity in viral DSPs (Figure 4B).

Our study highlights two ancient host-virus battles for Dynamin-like functions in eukaryotic lineages. First, we show that the Mx clade of DSPs is ancient and may represent an early DSP specialization for antiviral function in eukaryotic evolution. Focused studies on characterizing representatives of this clade in fungi, plants, and basal branching eukaryotes would reveal more insight into Mx function and mechanism. Second, we find a recurrent pattern of DSP acquisition by different lineages of NCLDVs, which also suggests that host and viral DSPs may perform critical yet poorly understudied functions, at least in this widespread lineage of viruses. Understanding how these putative arms races occur for cellular and viral membrane remodeling may reveal novel aspects of viral biology and host defense. Thus, besides their canonical cell biological roles in membrane remodeling, our study reveals that Dynamin-super-family proteins (DSPs) have played critical antiviral and viral functions through most of eukaryotic evolution.

## Supporting information

Accessions, alignments, and nexus tree files

## Acknowledgments

We thank members of the Malik and Emerman labs (especially Ching-Ho Chang, Will Henriques, and Janet Young) for valuable discussions and feedback on the manuscript. We also thank Ed Culbertson for their feedback on the manuscript. The authors would like to thank L. Aravind, whose observation of Mx-like sequences in algae spurred us to look more deeply at Mx evolution. This work was supported by the University of Washington Cellular and Molecular Biology Training Grant (T32 GM007270, to CAL), Howard Hughes Medical Institute Hanna H. Gray Fellowship (GT11096/GT16732 to JLT), American Foundation for AIDS Research Mathilde Krim Fellowship in Biomedical Research (110298-71-RKHF/110537-74-RKHF, to JLT), National Institutes of Health grant (U54 AI170792 (PI: Nevan Krogan) to ME, HSM), and a Howard Hughes Medical Institute Investigator award (to HSM). Funding agencies had no role to play in the execution of the project or the decision to publish. This paper was typeset with the bioRxiv word template by @Chrelli: www.github.com/chrelli/bioRxiv-word-template

## Author contributions

Conceptualization: CAL, PAD, ME, JLT, HSM

Methodology: CAL, PAD, HSM

Investigation: CAL, PAD, JLT, HSM

Visualization: CAL, PAD

Funding acquisition: JLT, ME, HSM

Project administration: HSM

Supervision: ME, HSM

Writing – original draft: CAL, PAD, HSM

Writing – review & editing: ME, JLT

## Materials and Methods

### Phylogenetic trees

DSPs were identified via iterative BLAST searches on representative species with fully sequenced genomes using *bona fide* DSPs as queries. In cases with multiple hits from the same species, only the longest isoform encoded by each DSP gene was utilized in downstream analysis. We used Clustal Omega (120) or MAFFT (55) to align sequences obtained from the same query. From these alignments, the GTPase domains were extracted and realigned using MAFFT to generate an alignment of GTPase domains from all DSPs. This alignment was then manually inspected to eliminate incomplete sequences and used to generate phylogenetic trees using FastTree (56, 57) with default settings, which was also used to perform bootstrap analyses. Trees were also subsequently built using IQ-Tree under default settings with UltraFast Bootstraps (58) using the LG+I+G4 model. Resulting trees were then manually annotated to indicate the level of bootstrap support and to highlight groupings. To analyze Dynamin Middle Domains and GEDs, sequences were aligned by Clustal Omega (120), and Dynamin Middle and GEDs were manually extracted and realigned. This alignment was then manually inspected to eliminate incomplete sequences and used to generate phylogenetic trees using FastTree (56, 57) with default settings, which was also used to perform bootstrap analyses. We expanded our search space in incremental steps of increasing phylogenetic coverage, starting first with animals, then to plants and fungi, then to all eukaryotes and viruses. More focused trees (Figure 1D, Figure 2C, Figure 3B and Figure 4) were excised from the larger trees (Figure 1A, Figure 2A, Figure 3A, respectively).

### Protein domain analysis

We uploaded full-length alignments of sequences retrieved from iterative Blast searches to the NCBI Conserved Domains tool using the Batch-CD search tool (60, 61). We present schematic versions of the domain analyses in our figures. We restrict our analyses only to those domains that were reliably identified by the NCBI CD tool. However, in certain cases, we additionally searched selected full-length DSP sequences using subsequent manual BLAST searches to confirm the absence of individual domains, such as the pleckstrin-homology (PH), Fuzzy onion (Fzo), or the TMP-synthase domains. We also visually inspected the GTPase domains identified to ensure that they bore all the hallmarks of catalytically active domains, including preservation of the previously identified catalytic residues.

### Cell biological localization

To represent the cytological location of different DSPs (Figure 2B), we focused on previously published cell biological studies in human cells.

### DSP repertoires in representative genomes

To comprehensively identify all DSPs in representative genomes, we performed BLAST searches to specific fully sequenced genomes using the ref_seq database using representative DSPs from each of the identified clades, taking care to correctly and comprehensively identify true paralogs. We used phylogenetic analyses and (when possible) shared synteny analyses to distinguish between orthologs and paralogs. These analyses allowed us to identify cases of gene duplication within clades and confidently identify cases of specific gene losses of specific DSP clades for each representative genome sequenced.

### GTPase catalytic residue alignment

Representative sequences were selected from each viral DSP clade. These were aligned, and the G1, G2/Signal I, G3, and G4 boxes were manually extracted and realigned using MAFFT. This alignment was run on ESPript 3 (107) with default settings to display the different G Box motifs.

**Supplementary Figure 1.**
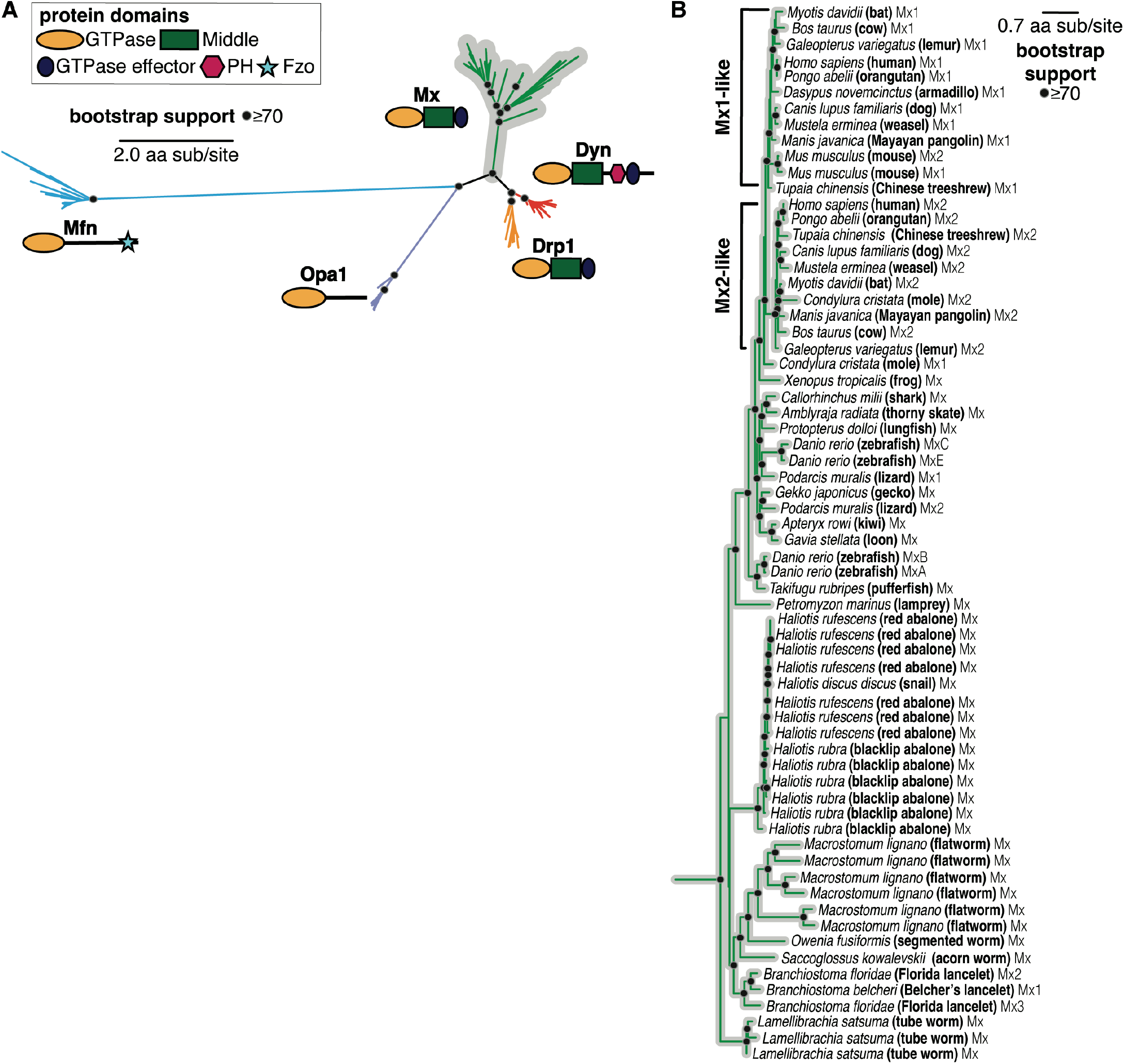
The evolutionary origin of Mx in animals predates the interferon signaling network. **(A**) Phylogenetic analysis of Dynamin-superfamily proteins (DSPs) based on their common GTPase domain in representative Holozoa (animals and their closest single-celled relatives) reveals five distinct DSP clades, consistent with Figure 1A. We used MAFFT to generate the alignment and IQ-Tree (112,113) to build the phylogeny. Black dots indicate nodes with UltraFast bootstrap support greater than 70% based on IQ-Tree analyses (see methods); a scale bar indicates the level of amino acid divergence. (B) Phylogenetic analysis of Mx-like proteins in animal species, consistent with Figure 1D.

**Supplementary Figure 2.**
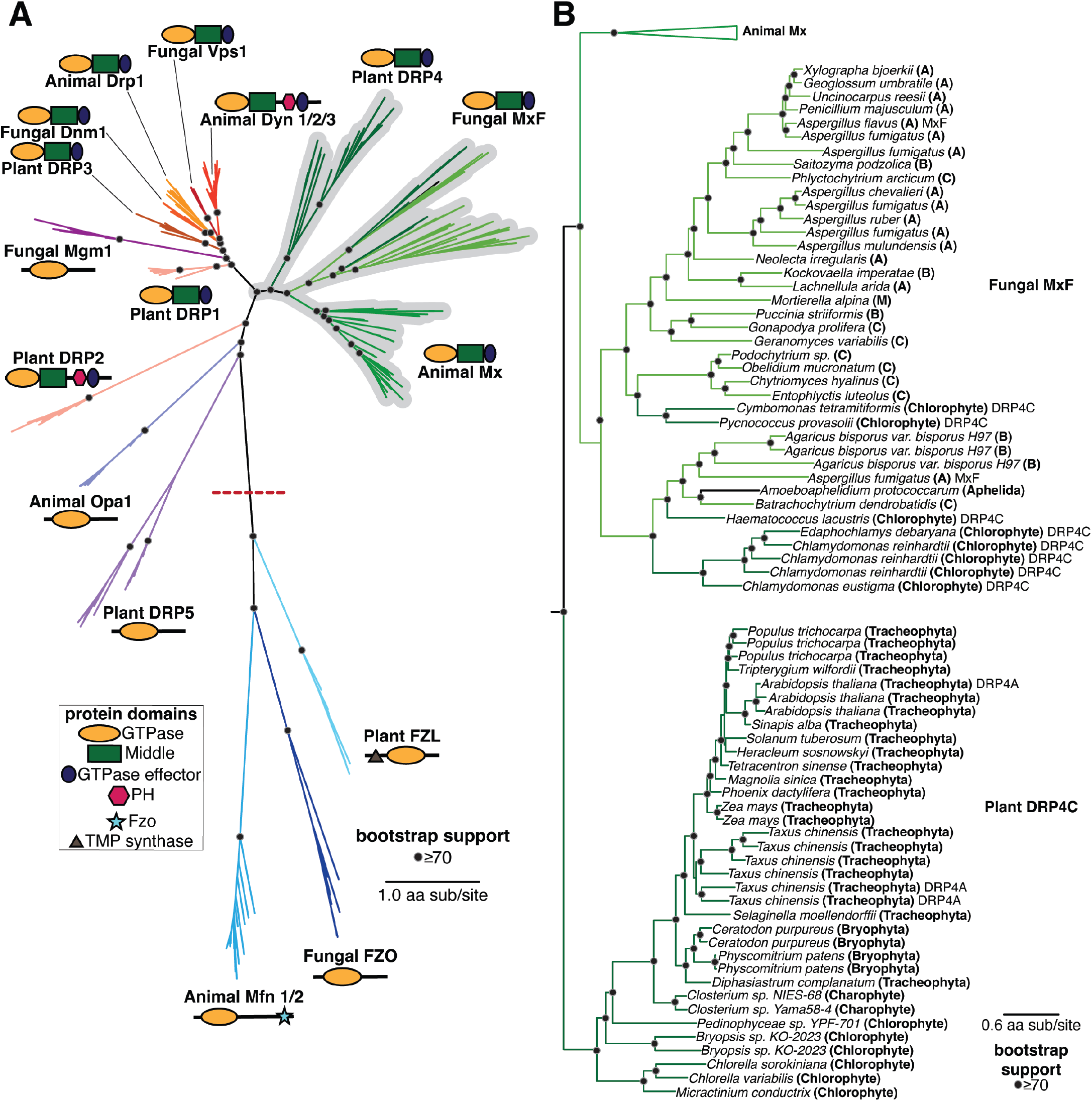
Animal DSP clades, including Mx proteins, in fungal and plant genomes. **(A)** Phylogenetic analysis of animal, fungal, and plant DSPs based on their common GTPase domain reveals broad groupings consistent with FastTree analysis in Figure 2A. We used MAFFT to generate the alignment and IQ-Tree (112,113) to build the phylogeny. Black dots indicate nodes with UltraFast bootstrap support greater than 70% based on IQ-Tree analyses (see methods); a scale bar indicates the level of amino acid divergence. **(B)** Phylogenetic analysis of animal Mx, fungal MxF (Mx-like proteins from Fungi), and plant DRP4 proteins, consistent with Figure 2C.

**Supplementary Figure 3.**
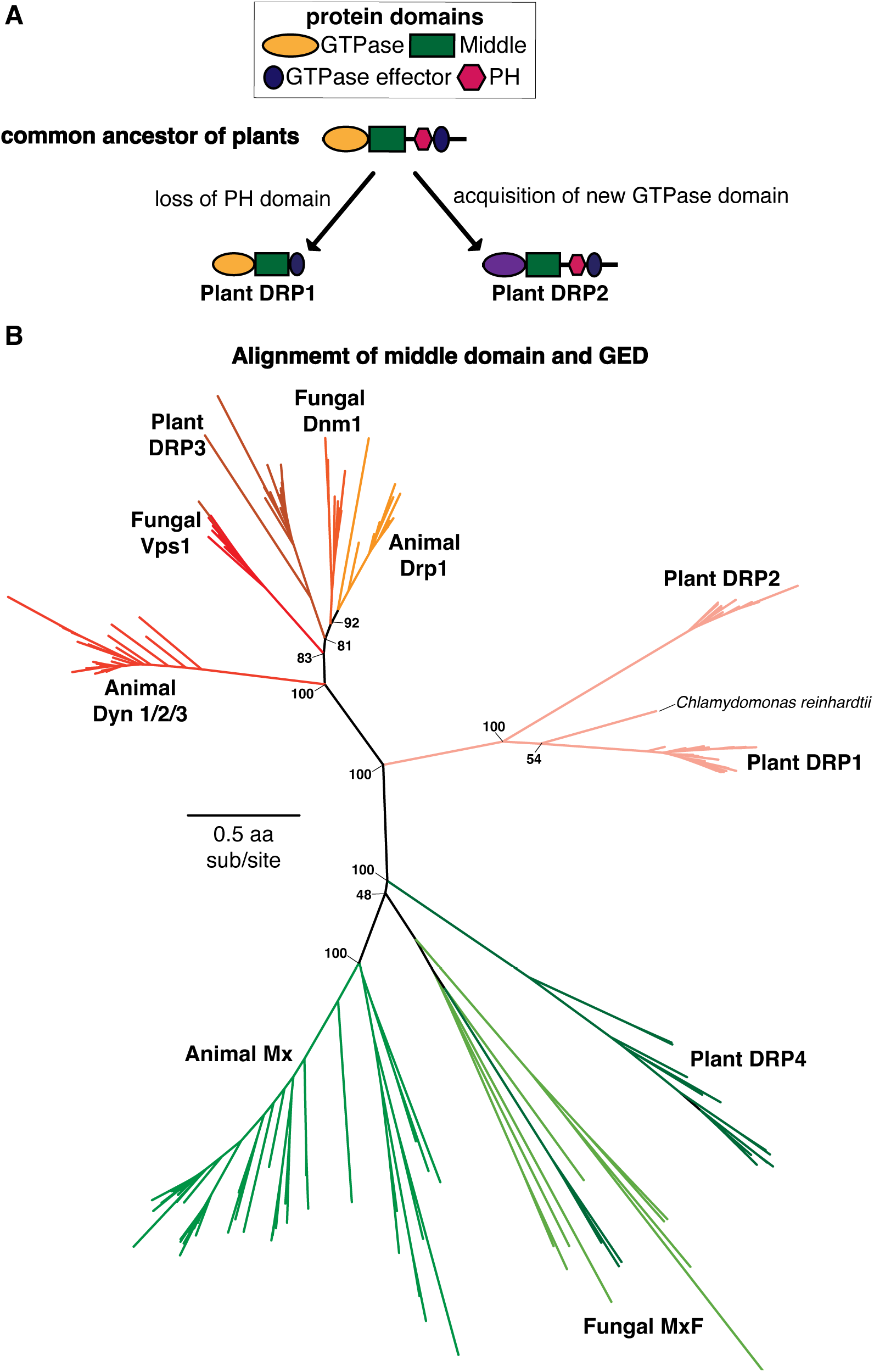
Evolutionary analyses of Plant DRP1 and DRP2 proteins. **(A)** A proposed model of Plant DRP1 and DRP2 evolution. The plant DRP1/DRP2 ancestor contained GTPase, Middle, GED, and PH domains. This ancestor was duplicated into DRP1, which lost its PH domains, and DRP2, which retained the PH domain but acquired a divergent GTPase domain via recombination. **(B)** Phylogenetic analysis of Plant, Animal, and Fungi DSP Middle and GED domains reveals that Plant DRP1 and DRP2 are much more closely related in the Middle-GED phylogeny than in the GTPase phylogeny (Figure 2). Although this does not clarify where the Plant DRP2 GTPase domain arose from (but see Figure 3), the rest of the protein appears to be derived from a common Plant DRP1/2 ancestor.

**Supplementary Figure 4.**
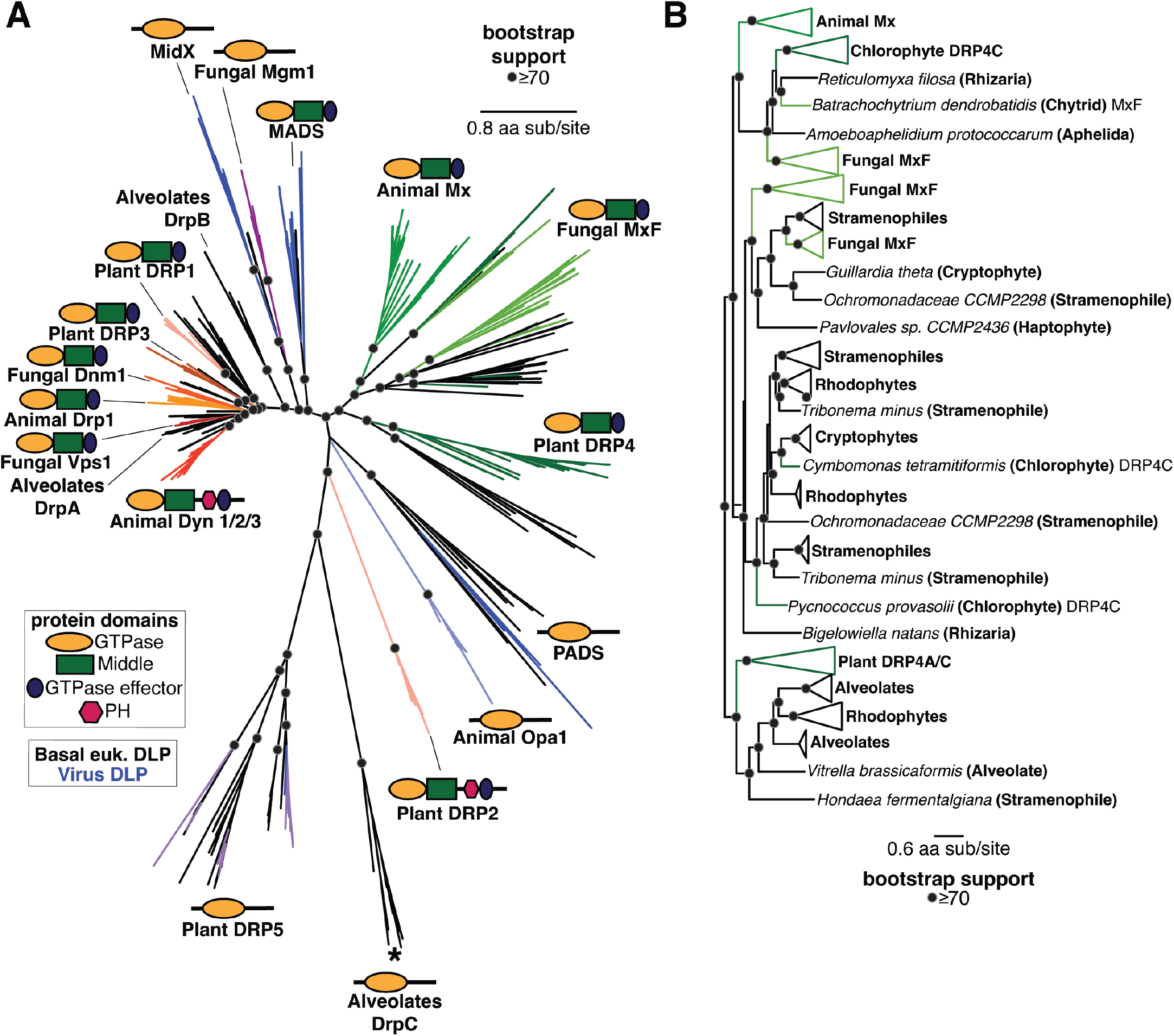
Phylogenetic analysis of DSPs in eukaryotes reveals an ancient Mx lineage. **(A)** Phylogenetic analysis of all eukaryotic DSPs (except for the Mfn/FZO, FZB, and FZL clades) reveals an ancient Dyn/Drp clade that includes Animal Drp, Plant DRP3, and Fungal Vsp1 along with fungal Dnm1, alveolate DrpA and DrpB, Plant DRP1, and DSPs from several basal branching eukaryotes (shown with black branches). The tree topology is consistent with FastTree analysis in Figure 3A. We used MAFFT to generate the alignment and IQ-Tree (112,113) to build the phylogeny. Black dots indicate nodes with UltraFast bootstrap support greater than 70% based on IQ-Tree analyses (see methods); a scale bar indicates the level of amino acid divergence. **(B)** Phylogenetic analysis of three deeply branching lineages of Mx-like proteins in eukaryotes, compared to Figure 3B

